# Environmental gradients and the establishment of chromosomal inversions

**DOI:** 10.64898/2026.10.25.761641

**Authors:** Stephan Peischl, Stefan Strütt

## Abstract

Chromosomal inversions are thought to promote local adaptation by capturing combinations of locally beneficial alleles and shielding them from recombination. Many of the best-documented examples are inversions that form clines along environmental gradients, yet analytical theory for this mechanism has remained confined to continent–island and two-deme models. We combine classical cline theory with quasi-linkage equilibrium approximations and spatial branching processes to derive the growth rate, establishment probability, and total establishment rate of a rare inversion along continuous environmental gradients of varying shape. After rescaling space by the characteristic cline width, all three quantities follow universal profiles and simple scaling laws. At a sharp environmental transition, the local growth rate of an inversion equals the selection acting on the captured alleles, independently of dispersal. Yet its establishment probability is only about one tenth of that value, again independently of dispersal, because a founding lineage disperses over the same spatial scale on which its fitness changes. In contrast to two-deme models, where the total establishment rate peaks at intermediate migration, along a cline it increases monotonically with dispersal, as successful inversions originate over an increasingly broad region on the flanks of the cline. Because the local growth rate depends only on the resident single-locus clines and the dispersal rate, the potential for inversion establishment can be estimated from observable allele-frequency clines without knowledge of the underlying fitness landscape. Stochastic simulations confirm these predictions. Our results provide a quantitative link between clinal patterns of genetic variation and the evolution of chromosomal inversions.

**Significance Statement:** Chromosomal inversions are thought to promote local adaptation by protecting co-adapted alleles from recombination, and many of the clearest examples form environmental clines. Yet for twenty years, theory for this mechanism has been limited to one or two discrete populations. We extend this theory to continuous environmental gradients. Two results differ qualitatively from discrete-population models. An inversion can grow as fast as selection on its captured alleles, yet its establishment probability is only about a tenth of that, regardless of dispersal. And whereas excessive gene flow hinders establishment in discrete models, along a cline the total establishment rate keeps increasing with dispersal. Gene flow, usually seen as an obstacle to local adaptation, can thus favour inversions that protect it.

## Introduction

Chromosomal inversions rearrange the order of a segment of a chromosome [1] and, in heterozygotes, strongly reduce recombination within the inverted region [2]. This suppression allows alleles captured by an inversion to be inherited as a largely non-recombining haplotype, while the same alleles continue to recombine on standard chromosomes. Inversions can therefore preserve combinations of co-adapted alleles that would otherwise be vulnerable to disruption by recombination [3, 4].

Inversions are frequently associated with local adaptation across diverse taxa: they segregate as geographic clines or occur at high frequency or near fixation in habitats experiencing contrasting selection [5–10]. Such associations have motivated the hypothesis that inversions promote local adaptation by protecting locally beneficial allele combinations from recombination and gene flow [11–16]. Notably, many of the best-known examples occur along environmental clines, where inversion frequencies change across continuous spatial gradients. For example, *In*(3R)*Payne* in *Drosophila melanogaster* has formed parallel latitudinal clines on multiple continents, consistent with local adaptation to spatially varying climatic conditions [17–21]. A similarly classic case is the 2La inversion in *Anopheles gambiae*, which varies repeatedly along aridity gradients in West and Central Africa: the inverted arrangement predominates in dry savannas, whereas the standard arrangement predominates in humid forests, a pattern consistent with selection for aridity tolerance [22, 23]. Recent work provides further examples. In Atlantic silversides, multiple large inversions show strong latitudinal clines along a steep thermal gradient and contribute in a modular fashion to continuous adaptive-trait variation despite extensive gene flow [24]. Likewise, in *Carex* on Réunion Island, large inversions concentrate genomic divergence across an island ecological gradient, consistent with a role in local adaptation [25].

Theory provides a mechanistic basis for these findings. Inversions can establish when they recover the recombination load that would otherwise erode co-adapted haplotypes [4, 26–31]. The foundational treatment by Kirkpatrick and Barton [4] proposed this mechanism in the context of a continuous-space model with two habitats. Their analytical treatment, however, considered a continent–island model, complemented by a stepping-stone simulation illustrating how the mechanism could operate in a spatially structured population. Subsequent analytical work has likewise focused on continent–island and two-deme models [26, 27, 32, 33]. These models have revealed that migration has a nontrivial effect on selection for inversions. In the continent–island model, the inversion growth rate, and hence establishment probability, increases approximately linearly with migration rate [4, 26]: migration introduces maladapted haplotypes and thereby creates a recombination load that favours reduced recombination. In two-deme models, however, this relationship is no longer monotonic. Although migration generates selection for inversions, sufficiently high migration can reduce their establishment probability by exposing inverted haplotypes to habitats in which they are not adapted [33]. Recent work has further incorporated mutational supply, capture of adapted haplotypes, and stochastic establishment to quantify total establishment rates [33].

More generally, modifier theory predicts that reduced recombination is favoured at migration–selection balance [34], although numerical and analytical studies of spatially structured populations have focused largely on conditions favouring increased recombination [35, 36]. Rear-rangements that tighten linkage among locally adapted loci can establish under gene flow in two-patch models [37, 38]. All of these studies, however, consider a small number of discrete populations.

Much less is known about how selection, recombination, and dispersal interact to favour or disfavour inversion establishment across continuous environmental gradients. This gap is somewhat surprising given the extensive theoretical framework for clines in continuous space, including well-established results for multilocus clines and the generation of linkage disequilibrium through spatial mixing [39–46]. In particular, the characteristic cline width *ω* is determined by the balance between dispersal and local selection [41, 42], while multilocus clines generate linkage disequilibrium through the mixing of differentiated gene pools [45, 46]. This spatial structure determines where recombination load accumulates and, consequently, where an inversion that suppresses recombination can be favoured. While continent–island and two-deme models can capture contrasting selection between discrete habitats, they do not describe how selection varies across an environmental transition or how cline width sets the spatial scale of linkage disequilibrium and recombination load. We therefore extend existing cline and QLE theory to the problem of inversion establishment in continuous space.

We analyse inversion establishment along a family of environmental gradients in continuous space. Using a two-locus diffusion framework and quasi-linkage equilibrium (QLE) approximations, we derive scaling laws for the inversion growth rate, establishment probability, and total establishment rate as functions of gradient shape, dispersal, and selection. Two results are especially relevant for both theoreticians and empiricists. First, total establishment rate behaves qualitatively differently from discrete two-deme predictions: in symmetric two-deme models it is typically maximized at intermediate migration [33], whereas in continuous space it increases monotonically with dispersal. Second, and perhaps more importantly, inversion establishment along a cline can be predicted from the resident single-locus clines and dispersal rates alone. After rescaling by cline width, the resulting scaling laws are independent of the exact spatial selection regime, meaning that the potential for inversion establishment can be inferred without detailed knowledge of the underlying fitness landscape.

## Model and Results

We consider a population distributed along a one-dimensional habitat where two bi-allelic loci, **A** and **B**, experience spatially varying selection. Let 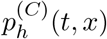 denote the frequency of haplotype *h* ∈ {*ab, aB, Ab, AB*} in chromosomal state *C* ∈ {*S, I*} (standard or inverted) at time *t* and position *x* ∈ R. The dynamics are

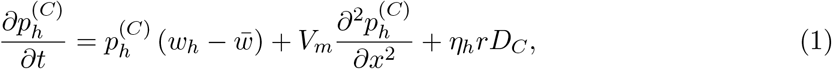

where *w*_*h*_ denotes the fitness of haplotype *h*, 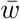 is the mean fitness, *V*_*m*_ is the dispersal variance, *r* is the recombination rate, *D*_*C*_ is linkage disequilibrium within chromosomal state *C*, and *η*_*h*_ = ∓1 for coupling (*AB, ab*) and repulsion (*aB, Ab*) haplotypes, respectively. We assume the population is initially fixed for the standard arrangement and that a stable two-locus cline has been established [47]. Hats (^) denote quantities evaluated at this resident equilibrium.

We ask for the local growth rate of a rare inversion that captures the locally adapted haplotype, say *AB*, when it is introduced into a resident population at equilibrium. Since establishment is determined by the dynamics of the inversion while it is rare, we linearize the system at this resident equilibrium and neglect the effect of the inversion on the resident haplotype frequencies. The frequency of the inverted *AB* haplotype then obeys

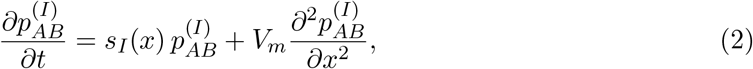

where 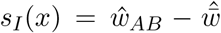 is the local invasion fitness, or per capita growth rate, of a rare inverted*AB* haplotype at position *x*. Using the resident equilibrium identity for 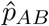 yields

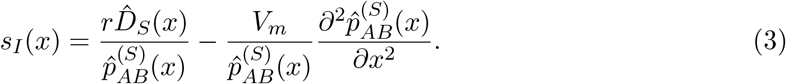

The first term represents recovery of the locally adapted haplotype through recombination load in the resident standard arrangement, whereas the second is a curvature term arising from migration–selection balance. Equation (3) is general: it does not assume quasi-linkage equilibrium (QLE) or a particular form of spatially varying selection, and it allows for dominance and epistasis.

If the inversion is introduced in proportion to equilibrium haplotype frequencies across space, the diffusion term in eq. (2) cancels the curvature contribution to *s*_*I*_ at the instant of introduction, so that the inversion frequency initially changes at the recombination-load rate 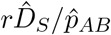 (Appendix A). This recovers classic results for the continent-island model: under weak migration with 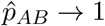 we get that a rare inversion increases at rate 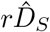, as previously obtained by Bürger and Akerman [27] and Charlesworth and Barton [26].

To evaluate eq. (3) we need exact two-locus cline solutions, which are generally unavailable. However, when recombination is fast relative to selection and to spatial turnover on the cline scale (*r* ≫ *s, r* ≫ *V*_*m*_*/ω*^2^)—the biologically relevant regime for inversion evolution, because otherwise suppressing recombination would add little— the QLE approximation allows us to express the inversion growth rate in terms of single-locus clines alone [47–49]:

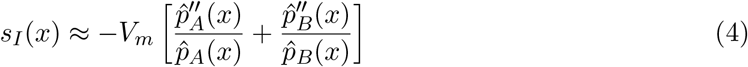

where 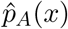 and 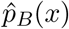 are the equilibrium frequencies of the locally adapted alleles *A* and *B*, respectively (Appendix B). This result is the foundation for our analysis; once a single-locus resident solution is known, establishment theory follows readily.

If dominance and epistasis are absent, as assumed throughout the remainder of this work, and *s*_*A*_(*x*) and *s*_*B*_(*x*) denote the local selection intensity favoring allele *A* and *B*, respectively, Equation (4) becomes

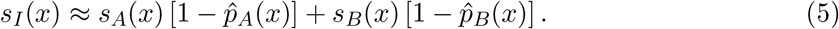

Each captured allele therefore contributes to inversion establishment in proportion to its local selective advantage and the local frequency of the maladapted allele, 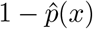, which reflects the migration load at that locus.

### Cline width scaling across environmental gradients

We next assume that selection is the same at both loci, and model spatially varying selection at each locus by *s*(*x*) = *s*_0_*g*(*x*), with the gradient family

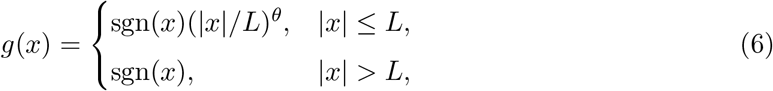

which interpolates from a sharp step (*θ* = 0) through gradual transitions (*θ* ≈ 1) to buffered profiles with peripheral ecotones (*θ* ≫ 1). Here *L* sets the spatial breadth of the transition and *θ* its shape (Fig. 1).

**Figure 1.**
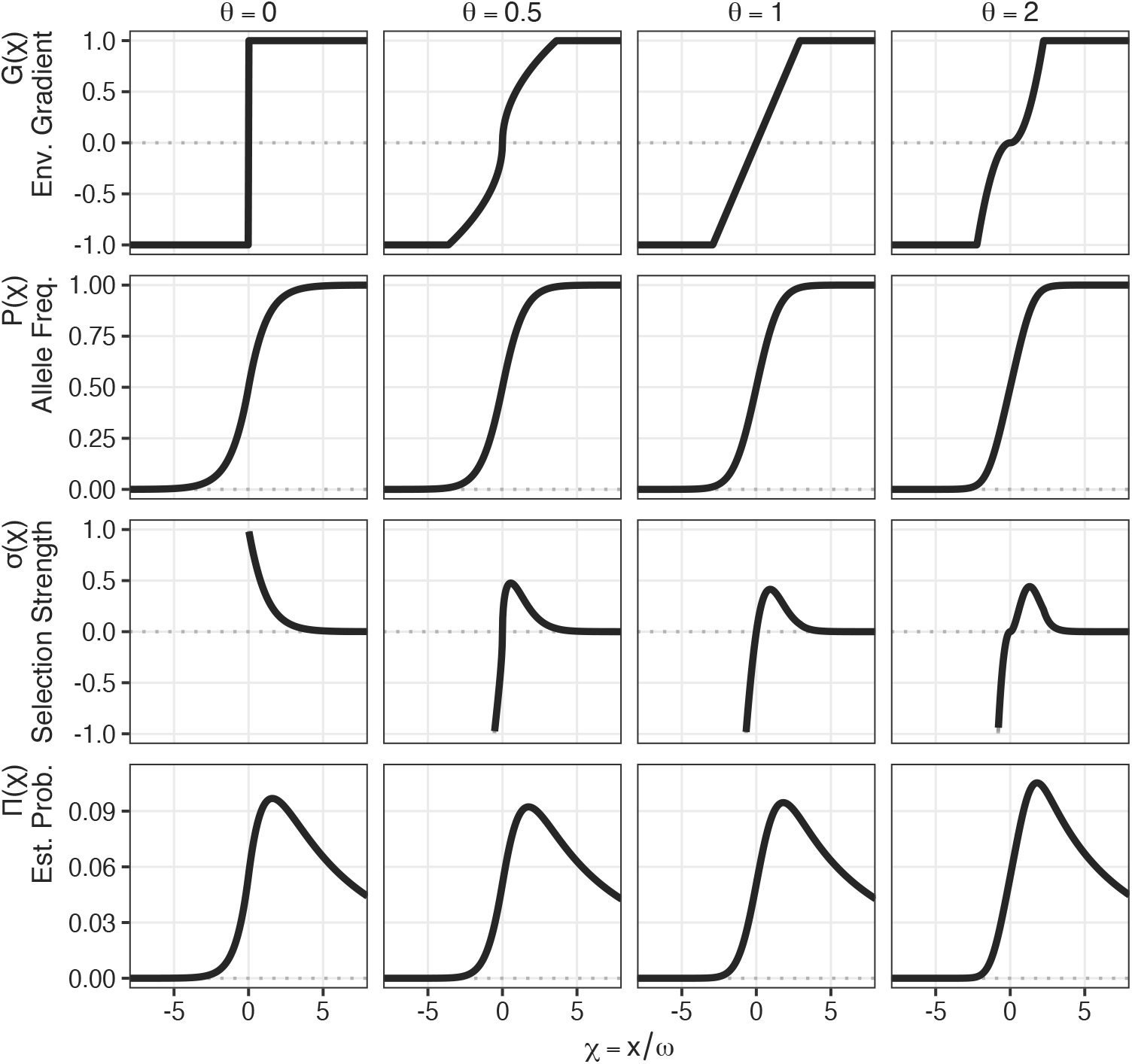
Spatial profiles after rescaling by cline width *ω*. Rows show the environmental gradient *G*(*χ*), the single-locus cline *P* (*χ*), dimensionless inversion growth rate *σ*(*χ*), and establishment probability Π(*χ*); columns vary gradient shape *θ*. The environmental transition width is 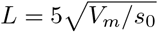.

Allele-frequency clines have a characteristic width *ω*, set by the balance between dispersal and local selection. A single-locus cline *P* (*χ*) in the rescaled coordinate *χ* = *x/ω* satisfies

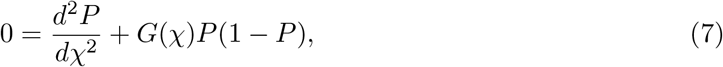

where *G*(*χ*) = *g*(*ωχ*) and *P* (*χ*) = *p*(*ωχ*). Choosing

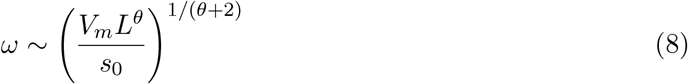

removes explicit dependence on *V*_*m*_ and *s*_0_ within each gradient class (Appendix D). The classical step scaling 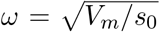 is recovered at *θ* = 0 [41], and the linear-gradient scaling *ω* = (*V*_*m*_*L/s*_0_)^1*/*3^ at *θ* = 1 [42]. Fig. 1 shows the resulting environmental and allele frequency profiles.

### Local inversion growth rate scales as *V*_*m*_*/ω*^2^

Under QLE we can write (Appendix C),

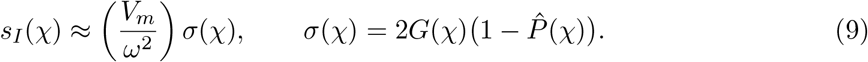

Here *σ*(*χ*) is the dimensionless growth-rate profile in the scaled coordinate *χ* = *x/ω*: after the magnitude *V*_*m*_*/ω*^2^ is factored out, the remaining spatial shape depends only on the rescaled environmental gradient *G*(*χ*) and the resident cline 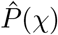. Thus (Appendix C)

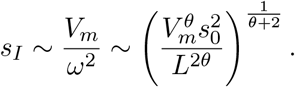

In a step environment the maximum of *s*_*I*_ is attained right at the border between the two habitats, where 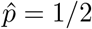, and equals *s*_0_: the inversion growth rate is then as large as selection on the captured alleles, and independent of dispersal rates. This reveals an interesting contrast to discrete models, where the inversion growth rate depends strongly on migration rates [26, 33]. For *θ >* 0 the local growth rate is lower, bounded by order *s*_0_(*ω/L*)^*θ*^, and displaced toward the periphery of the cline (Fig. 1). When the environmental gradient is flatter (*θ >* 0), dispersal becomes progressively more important as transitions flatten. For instance, for *θ* → ∞ the inversion growth rate scales with *V*_*m*_*/L*^2^.

Note that if *L < ω*, rescaling of space breaks down and the system behaves like a step. When *L > ω*, increasing *θ* reduces the maximum of *s*_*I*_ while broadening the region of positive growth. Exact two-locus PDE solutions confirm the QLE profiles across gradient shapes (Fig. 2). For *n* ≥ 2 loosely linked loci under QLE, growth rates add across captured clines (Appendix G),

**Figure 2.**
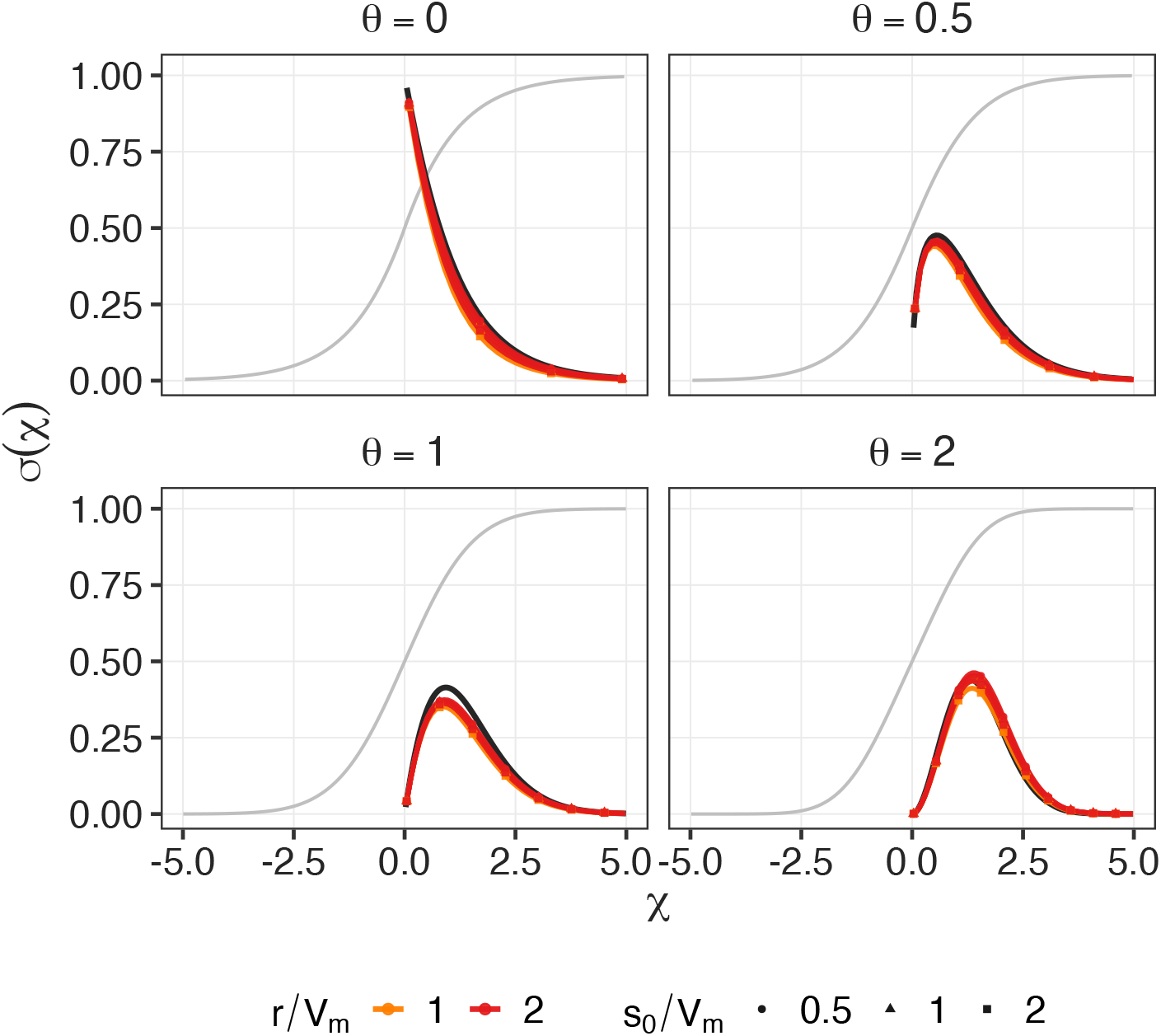
Spatial profile of the inversion growth rate *s*_*I*_. The gray line shows the equilibrium single locus allele frequency clines. The solid black line shows the QLE prediction from eq. (9), using these cline solutions. Colored lines show exact numerical solutions of the full two-locus PDE across various parameter combinations. After rescaling, the numerical solutions become virtually indistinguishable from each other, and follow the QLE approximation very closely. In the left half of the habitat, inversions are negatively selected, but the y-axis is truncated at zero for visual clarity.

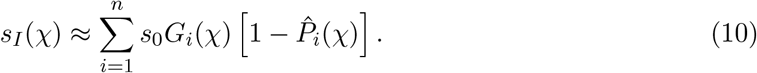

### Establishment probability of inversions

The establishment probability *π*(*x*) of a rare inversion can be approximated by a continuous-space branching-process equation [50, 51]. Rescaling yields the parameter-free problem

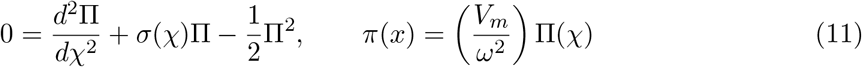

(Appendix E). For a given environmental gradient, the shape of Π(*χ*) is therefore universal up to scaling (Fig. 1), and the inversion establishment probability scales with the local growth rate of inversions.

With spatially uniform selection, establishment probability is typically around twice the selective advantage of a mutation [52]. At a step cline centre, *s*_*I*_ = *s*_0_, thus one would assume that the establishment probability should be close to that if dispersal is limited. Yet we find that Π_max_ ≈ 0.1 and thus *π*_max_ ~ *s*_0_*/*10, independently of how strong dispersal is. This counterintuitive result can be explained by observing that even though the magnitude of *s*_*I*_ does not depend on dispersal, the spatial range of positive growth is driven by dispersal itself. The region with strongly positive growth is set by migration–selection balance (*ω*), and dispersal occurs on the same spatial scale as selection for the inversion. Thus, independently of how strong dispersal is, an inverted lineage occurring at a given location *x* will always experience the same selective environment over the course of its establishment.

We further find that Π(*χ*) peaks away from the centre, even when *s*_*I*_ is maximal right at the center (step environment, *θ* = 0). This is because the growth-rate profile *σ*(*χ*) is asymmetric around the center of the cline. On one side, *s*_*I*_ falls steeply into deleterious habitat; on the other, it declines more slowly into a long near-neutral tail. Stochastic Wright–Fisher simulations of a stepping-stone model show that the branching-process approximation is accurate (Fig. 3).

**Figure 3.**
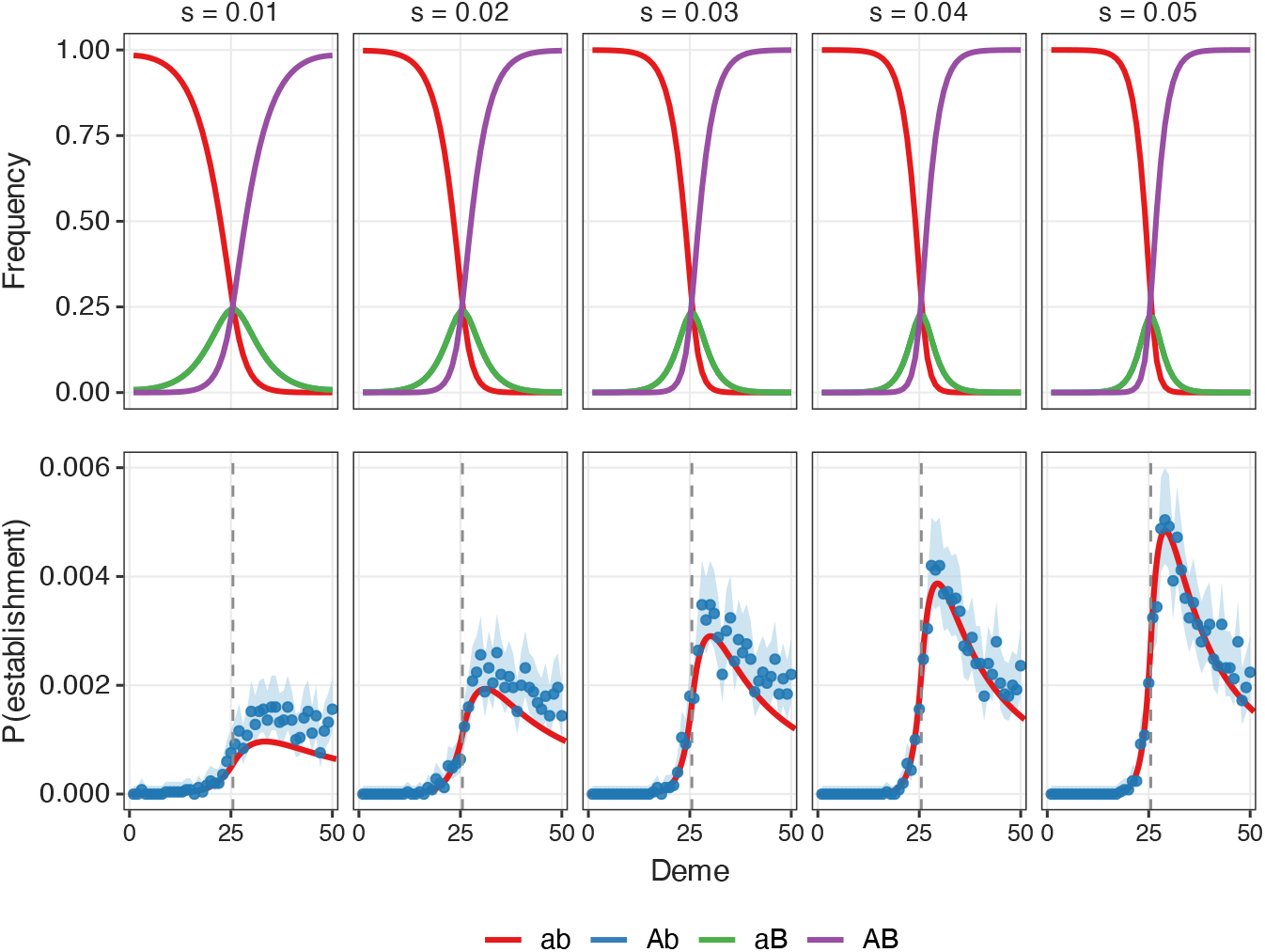
Wright–Fisher stepping-stone validation of establishment probabilities. Top: equilibrium haplotype frequencies after burn-in under a stepped environment (demes 1–25 vs. 26–50). Bottom: establishment probability versus deme of introduction (points with Wilson 95% CIs) compared with the continuous-space branching-process prediction (curves). Simulations use 50 demes with *N* = 2000 individuals each, migration rate *m* = 0.5, recombination rate *r* = 0.5, and selection coefficients *s* ∈ {0.01, …, 0.05} (25,000 replicates per deme). A rare inversion is introduced as a single inverted *AB* copy after a 2000-generation burn-in and scored as established if its global frequency exceeds 0.05 (i.e., 5% of the total population) within 200,000 generations; lineages still present but below threshold at the time limit count as failed. Theory curves solve the same step-environment branching-process equation as in Fig. 1, with dispersal variance *V*_*m*_ = *m/*2 matching the stepping-stone migration scheme and continuous position mapped to deme coordinate as deme = (50 + 1)*/*2 + *x*.

### Total establishment rate

The total establishment rate across the habitat depends on both the spatial variation in establishment probability and the availability of mutational targets. Locally adapted haplotypes, which provide the mutational targets for inversions, are more common towards the periphery of the cline, further shifting the origin of successful inversions away from the center. Our results therefore suggest that successful inversions are expected to arise over a broad geographic range, while they will reach high frequencies near the habitat center where selection is strongest. Individual-based SLiM simulations [53] on a stepped 20-deme habitat confirm this pattern for both two-locus and four-locus architectures (Fig. 4).

**Figure 4.**
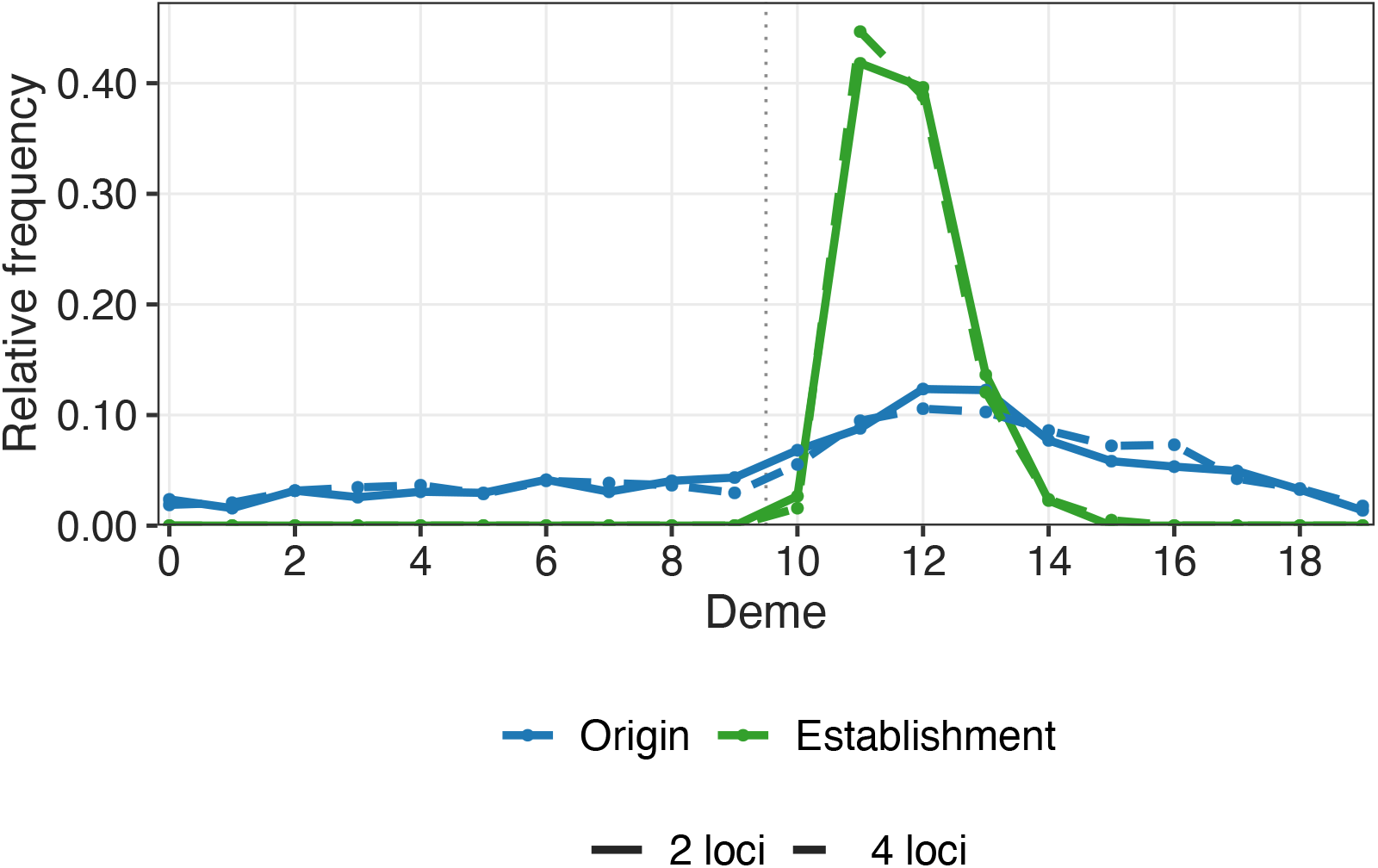
Origin and establishment locations of successful inversions. For each successful inversion, we show the distribution of the deme in which it first arose (*Origin*, blue) and the deme in which its frequency first exceeded the establishment threshold (*Establishment*, green), for two-locus (solid) and four-locus (dash-dotted) architectures obtained from individual-based, forward-in-time SLiM simulations. The dotted vertical line marks the habitat boundary (deme 9.5). Selection coefficients were calibrated so that a fully locally maladapted haplotype had the same relative fitness under both architectures: (1 − 0.2)^2^ = 0.64 for the two-locus model and (1 − 0.106)^4^ ≈ 0.64 for the four-locus model. Established inversions carried the locally adapted haplotype appropriate to the habitat in which they established. For clarity, only established inversions carrying the right-adapted haplotype (*AB* or *ABCD*, favored in demes 10–19) are shown. Histogram densities are normalized within each distribution (*n* = 2,000 established inversions per scenario).

We can gain further insights by integrating the local establishment probability against mutational supply, which gives the total rate at which successful inversions enter the population,

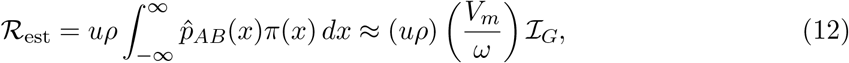

where *u* is the inversion mutation rate, *ρ* is population density, and ℒ_*G*_ is a dimensionless integral determined by the geometry of the environmental gradient (Appendix F). Thus ℛ_*est*_ ~ *V*_*m*_*/ω*.

More explicitly,

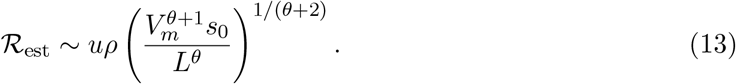

For a step environment, 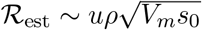, indicating a symmetric dependence on dispersal and selection. Both the local selective advantage *s*_*I*_ and the establishment probability *π* scale with *s*_0_, whereas the geographic range of establishment widens with dispersal as 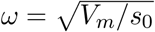. Thus, the 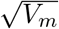 dependence of ℛ_est_ reflects an expansion of the region in which inversions can originate and establish, rather than an increase in their local probability of establishment. For a linear gradient the scaling becomes 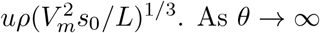,

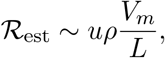

and the rate is controlled by dispersal and the geographic scale *L*, with little sensitivity to *s*_0_. Wright–Fisher stepping-stone simulations confirm this scaling: estimated total establishment rates collapse onto the predicted power law across dispersal, selection, and gradient shape (Fig. 5).

**Figure 5.**
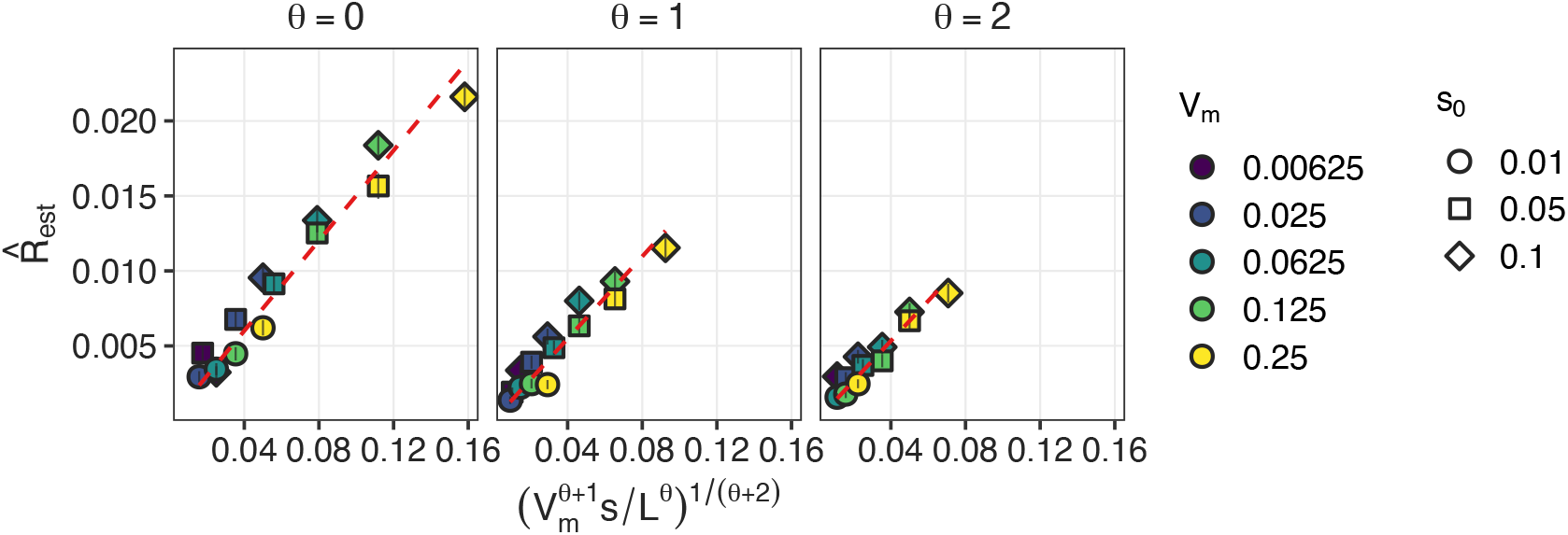
Validation of the total establishment-rate scaling (eq. (13)). Points: Wright–Fisher estimates 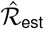 from 10^5^ single-origin establishment assays per parameter combination (50 demes, *N* = 2000, *r* = 0.5, *m* ∈ {0.0125, 0.05, 0.125, 0.25, 0.5}, *s*_0_ ∈ {0.01, 0.05, 0.1}, *θ* ∈ {0, 1, 2}, 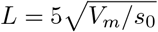, global establishment threshold 5% of total population). Vertical bars: standard errors from binomial sampling of 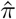. Dashed line: ordinary least-squares fit through the origin within each panel. Dispersal variance was chosen as *V*_*m*_ = *m/*2, where *m* is the stepping-stone nearest-neighbour migration rate. The horizontal axis is the theoretical scale 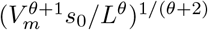.

## Discussion

We have extended classical cline theory [41, 45] to the establishment of rare chromosomal inversions along continuous environmental gradients. Across a broad range of environmental transition shapes and widths, rescaling space by the characteristic cline width reveals universal patterns of allele frequencies, inversion growth, and establishment (Fig. 1). The resulting scaling laws provide a simple description of how the spatial scale of adaptation determines the inversion growth rate: the magnitude of *s*_*I*_ scales with *V*_*m*_*/ω*^2^ (Appendix C), while its spatial profile is invariant after rescaling space by *ω*. When integrated across the habitat, this spatial scaling leads to a total establishment rate proportional to *V*_*m*_*/ω*.

Our analysis recovers and extends classical discrete-space results. When the inverted haplotype is introduced proportionally to its standard counterpart—for example, by replacing a fixed proportion of standard $AB$ haplotypes with inverted $AB$ haplotypes—the inversion frequency initially changes at the recombination-load rate 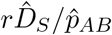. This recovers the exact result of Charlesworth and Barton [26], which we express in a form that makes the connection to recombination load explicit. Under weak migration, 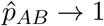, so that 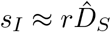, recovering the result noted by Charlesworth and Barton [26], Bürger and Akerman [27].

The growth rate of rare inversions can be very strong, matching the strength of selection at the loci they capture (eq. (9)). However, the establishment probability of an initially rare inversion remains roughly 20-fold lower than the no-dispersal expectation of *π* ≈ 2*s* (Fig. 1; cf. Fig. 3). The reason is that, even when *s*_*I*_ is independent of dispersal rates, dispersal determines both the movement of lineages across space and the spatial extent over which positive or negative growth occurs. The growth-rate profiles derived here show why this is costly: the deleterious effect of an inversion in the wrong environment is substantially stronger than its growth rate in the favored habitat (Fig. 1, *σ* row). Wright–Fisher simulations confirm this branching-process prediction for the establishment probability *π*(*x*) (Fig. 3).

Importantly, this does not imply that establishment is confined to a narrow region: dispersal broadens the spatial range over which inversions can establish, so locally adapted inversions may arise over a broad geographic range even though those that establish are favored near the habitat center (Fig. 4). Thus, the spatial locations where *s*_*I*_ is largest are not necessarily those that contribute most to its establishment (Figs. 3 and 4), highlighting an important consequence of spatially continuous selection that is absent from simpler discrete-population models.

When considering the total establishment rate across the whole habitat, averaging over the mutational target of the inversion, we find notable differences between discrete and continuous models. In the continuous-space model, the total establishment rate ℛ_est_ increases with both migration and dispersal. In contrast, in two-deme models, total establishment is maximized at intermediate migration rates. Too little migration limits the supply of maladapted haplotypes that generate selection for inversions, whereas too much migration homogenizes the habitat, exposes inversions to strongly deleterious backgrounds, and reduces their mutational target [33]. In one-dimensional continuous space, the negative effect of large dispersal is buffered: lineages originating sufficiently far from the center can still experience a predominantly favorable background and establish (Figs. 3 and 4). Moreover, the locally adapted haplotypes that form the mutational target of inversions are most abundant away from the center. Consequently, these regions can contribute substantially to total establishment despite having a weaker local growth rate *s*_*I*_ than the habitat center.

These results yield two clear takeaways for empirical research. First, continuous and discrete models make different predictions about how gene flow affects the fate of inversions, with strong dispersal monotonically increasing the rate at which inversions establish in clinal models. Second, and we regard this as the central practical message, inversion dynamics along a cline can be understood from resident single-locus cline shape and dispersal alone. Under QLE, the local inversion growth rate and establishment probability follow from the equilibrium cline 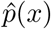 and *V*_*m*_. Mapping an allele-frequency cline and estimating dispersal rates therefore yields the main ingredients needed to apply the theory.

Several assumptions underlying our analysis delimit its current scope. We assumed complete suppression of recombination in heterozygotes, no epistasis, no dominance or underdominance between arrangements or alleles, and a one-dimensional, spatially symmetric environment. These assumptions keep the analysis tractable but are not fundamental to the QLE framework used here [41, 45, 47]. Qualitatively, underdominance between chromosomal arrangements is expected to reduce the inversion growth rate, whereas positive epistasis among captured alleles should increase it. The effect of dominance is more difficult to predict a priori, as it can alter the shape and symmetry of the resident cline and thereby modify *s*_*I*_. Because eq. (3) separates the recovery of recombination load from the spatial structure of the resident cline, the framework should extend to partial, cis-acting suppressors of recombination, such as other rearrangements or local changes in crossover rate, whereas loosely linked modifiers experience only weaker indirect selection [36]. We have also focused primarily on a biallelic, two-locus system and assumed that recombination is fast relative to selection and spatial turnover on the cline scale (*r* ≫ *s*_0_, *r* ≫ *V*_*m*_*/ω*^2^). Under this QLE regime, the effects of individual loci add approximately independently (Appendix G), such that tightly linked loci can be treated as a single selected block, whereas more loosely linked blocks contribute approximately additively. When selection becomes stronger relative to recombination, this approximation breaks down because selection maintains linkage disequilibrium on the standard background, reducing the recombination load that an inversion can recover and hence the growth rate gained by suppressing recombination [33]. These considerations suggest that inversions should be most strongly favored by genetic architectures comprising many loci of small to intermediate effect, separated by appreciable recombination distances, rather than by a few loci of large effect. This may also help explain why targets of selection within inversions can be difficult to identify from divergence between arrangements: many loci of small effect should produce less pronounced divergence patterns than a few loci under stronger selection [54].

A major question in the study of inversions and local adaptation is whether inversions arose by capturing locally adapted haplotypes [4], or through alternative routes to establishment, including drift, direct fitness effects of the rearrangement, meiotic drive [12], or associative overdominance from sheltered load [16]. Our results do not provide direct evidence for or against these hypotheses. They do, however, clarify the conditions under which capture is expected to be effective: sharp environmental transitions and strong dispersal. Sharp transitions increase the inversion growth rate *s*_*I*_, while greater dispersal increases the total rate of successful establishment across the habitat. This may help explain the prevalence of inversion polymorphisms along geographically extensive clines.

## Methods

### Wright–Fisher stepping-stone simulations

To test the continuous-space branching-process predictions, we simulated a one-dimensional stepping-stone model under Wright–Fisher sampling. The habitat consisted of a linear array of demes with nearest-neighbour migration. Within each deme, a fixed number of diploid individuals were formed by multinomial sampling from expected haplotype frequencies after selection, recombination, and migration. The two selected loci segregated on a standard (non-inverted) chromosomal background together with an inverted background that suppressed recombination between the captured loci. Fitness was multiplicative across loci, with locally adapted alleles favoured on opposite sides of a step (or power-law) environmental transition.

Dispersal was implemented as a discrete Laplacian on the deme lattice: interior demes exchanged migrants at rate *m/*2 with each neighbour, corresponding to continuous-space dispersal variance *V*_*m*_ = *m/*2. Boundary demes retained a reflecting migration scheme. Recombination among standard haplotypes occurred at rate *r*; inverted haplotypes did not recombine between the captured loci.

In both protocols below, each parameter set began with a deterministic burn-in: starting from equal frequencies of the four standard haplotypes and zero inversion frequency, we iterated the difference equations until an equilibrium without inversions is reached. The resulting standard-arrangement cline was then used as the initial condition for independent stochastic establishment assays.

#### Establishment probability profiles (Fig. 3)

We used 50 demes with *N* = 2000 individuals each, migration rate *m* = 0.5, recombination rate *r* = 0.5, and selection coefficients *s* ∈ {0.01, …, 0.05} under a stepped environment (demes 1–25 vs. 26–50). After a 2000-generation deterministic burn-in for each *s*, each assay copied that equilibrium, introduced a single inverted *AB* haplotype into a chosen deme (frequency 1*/N*, reducing the local standard haplotypes accordingly), and then evolved under full Wright–Fisher sampling with no further inversion mutation. The inversion was scored as established if its global frequency exceeded 5% of the total population within 200,000 generations, and as extinct otherwise; lineages still present but below threshold at the time limit were not counted as established. We ran 25,000 independent introductions per deme and compared the resulting establishment probabilities (with Wilson 95% confidence intervals) to numerical solutions of the continuous branching-process equation for the same step environment, mapping continuous position to deme coordinate as deme = (50+1)*/*2+*x*. Both Wright–Fisher protocols use the same strict global establishment criterion as Fig. 5.

#### Total establishment rate (Fig. 5)

To validate the scaling 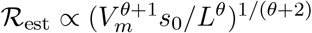, we used the same lattice size and deme size (50 demes, *N* = 2000) with *r* = 0.5, environmental transition width 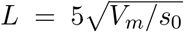, and a global establishment threshold of 5% of the total population. We varied nearest-neighbour migration rate *m* ∈ {0.0125, 0.05, 0.125, 0.25, 0.5} (with *V*_*m*_ = *m/*2), selection *s*_0_ ∈ {0.01, 0.05, 0.1}, and gradient shape *θ* ∈ {0, 1, 2}. After a 3000-generation deterministic burn-in for each (*m, s*_0_, *θ*) combination, we estimated the rate at which inversions establish by tracking single introductions: the introduction deme was drawn in proportion to the local equilibrium frequency of the locally adapted haplotype 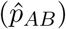, one inverted copy was inserted, and we recorded whether it established. Repeating this 10^5^ times per parameter combination gives the fraction of successful introductions, or equivalently the mean number of introductions needed until one establishes. Assuming that inversions arise so rarely that concurrent mutants do not interfere with each other, multiplying this success rate with the size of the mutational target (the number of adapted haplotype copies across the habitat) yields an estimate of ℛ_est_ up to the exact value of the mutation rate. Estimated rates were plotted against the predicted theoretical scale 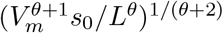.

### Individual-based SLiM simulations

Geographic origins and establishment sites of successful inversions (Fig. 4) were obtained from individual-based, forward-in-time simulations in SLiM 5 [53, 55]. Simulations comprised a linear stepping-stone array of 20 demes with total population size *N* = 5,000, split evenly across demes, and nearest-neighbour migration (*m* = 0.25). In the two boundary demes, inversion-bearing haplosomes were inviable (relative fitness =0), so that these demes acted as sinks preventing spurious edge effects of accumulating inversions. Two adaptive loci were positioned at relative positions 0.1 and 0.9 within a 1 Mb inversion on a chromosome with recombination rate *r* = 10^*−*7^ per bp per generation. In the four-locus architecture, two additional loci were added at relative positions 0.37 and 0.63, dividing the interval between the outer loci into three equal segments, under the same recombination rate. Alleles *a* and *b* (and *c* and *d* in the four-locus model) were favoured in demes 0–9, whereas alleles *A* and *B* (and *C* and *D*) were favoured in demes 10–19. All simulation parameters were identical between the two architectures except the selection coefficient, which was calibrated so that a haplotype fully mismatched with the local habitat had the same relative fitness under both architectures. The per-locus selection coefficient was *s* = 0.2 in the two-locus model, giving (1 − *s*)^2^ = 0.64, and 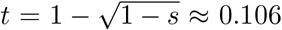 in the four-locus model, giving (1 − *t*)^4^ = 0.64. Newly arising inversions occurred at rate 10^*−*4^ per haplosome per generation after a 1,000-generation burn-in. Inversions arose irrespective of genotype at the adaptive loci and therefore captured the haplotype present on their chromosome of origin. Recombination between inverted and standard arrangements was fully suppressed. An inversion was considered established once its frequency exceeded 0.2 in any non-boundary deme. We ran 2,000 replicates per scenario.

## Code Availability

Simulation code and figure scripts are available at https://github.com/s-peischl/InversionFigures.

## AI use statement

AI-based tools were used in a limited supporting capacity for language editing and proofreading, for improving code documentation and reproducibility, for preparing code and associated materials for public release on GitHub, and for suggestions concerning figure design and presentation. All scientific analyses, mathematical derivations, computational methods and coding, as well as the initial preparation of the manuscript, were conducted by the authors without the use of AI tools. All AI-assisted outputs were critically reviewed and revised by the authors, who take full responsibility for the content and conclusions of the manuscript.

## Acknowledgments

We thank Claudia Bank and Julio Antonio Ayala Lopez for feedback on an earlier version of this manuscript. We also thank the members of the THEE and ComBi groups for feedback during lab meetings. SS and SP were supported by the Swiss National Science Foundation project grant 10001034 (Evolutionary dynamics of chromosomal inversions and local adaptation) awarded to SP.

## Appendix

**Table 1.** Model Parameters and Quantities.

| Symbol | Parameter / Quantity | Description / Context |
| --- | --- | --- |
| $V_m$ | Diffusion coefficient | Variance of diffusive individual dispersal |
| $r$ | Recombination rate | Rate of recombination in the standard arrangement |
| $s_0$ | Selection intensity | Strength of selection on locally adapted alleles (maximum over the spatial transition) |
| $L$ | Transition length | Spatial length scale of the selection transition |
| $\theta$ | Shape parameter | Controls the geometry of the selective transition |
| $\omega$ | Characteristic cline width | Scale of spatial variation in allele frequencies |
| $x$ | Position | Physical spatial coordinate |
| $\chi$ | Dimensionless position | Scaled spatial position, $\chi = x/\omega$ |
| $s_I(x)$ | Local inversion growth rate | Spatial profile of the inversion growth rate $s_I = w_{AB} - \bar{w}$ |
| $\sigma(\chi)$ | Dimensionless inversion growth rate | Scaled profile $s_I = (V_m/\omega^2) \sigma(\chi)$ |
| $\pi(x)$ | Establishment probability | Spatial profile of the inversion's establishment probability |
| $\Pi(\chi)$ | Dimensionless establishment | Scaled profile of the inversion establishment probability |
| $\mathcal{R}_{est}$ | Total successful input rate | Overall rate of successful inversion establishment |

### Appendix A: General Formula for the Inversion Growth Rate

We consider a spatially localized introduction of an inversion capturing the locally adapted haplotype *AB*. The resident population is initially at equilibrium 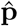 and fixed for the standard arrangement (*S*). The inversion is introduced at absolute frequency *ϕ*(*x*), so that at *t* = 0

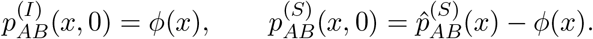

All other haplotype frequencies remain at their equilibrium values. Because the inversion is a neutral recombination modifier, this perturbation does not change haplotype fitnesses *w*_*h*_ or mean fitness 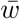 to linear order in *ϕ*.

While the inversion remains rare, recombination on the inverted background can be neglected, and eq. (1) reduces to the PDE for the inverted haplotype,

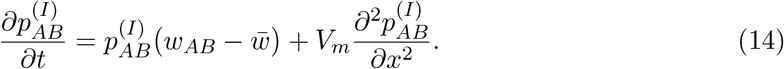

We write 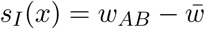 for the local growth rate in this PDE, so that

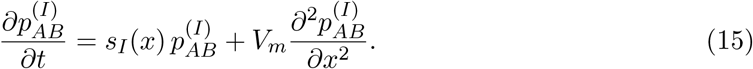

The first term is local growth at rate *s*_*I*_; the second is diffusion. Both remain present throughout the rare-inversion dynamics.

The growth rate *s*_*I*_ is fixed by the resident equilibrium. For the standard *AB* haplotype,

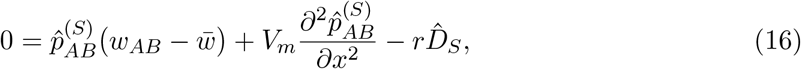

and therefore

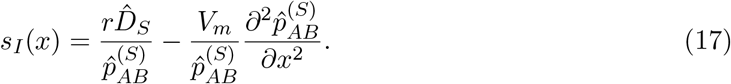

This is eq. (3) of the main text. The first term is recombination-load recovery: the inversion suppresses recombination and therefore gains the local disequilibrium that was breaking up the adapted haplotype. The second term is the curvature contribution required by migration–selection balance of the resident haplotype.

At the instant of introduction, we set 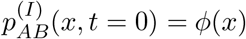, where *ϕ*(*x*) describes the density of the inversion right after it appears. The relative rate of change of inversion frequency is

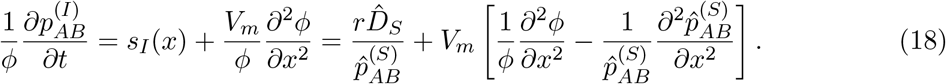

If the inversion is introduced in proportion to the resident *AB* frequency, 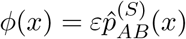 for small *ε*, the two curvature terms cancel and

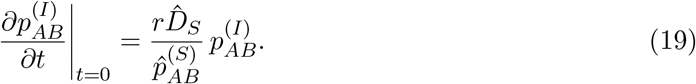

Thus, immediately after a proportional introduction, the inversion frequency changes at the local recombination-load rate 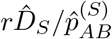. This coincides with the instantaneous growth rate derived for continent-island models [26]. Under weak migration with 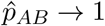, this further reduces to 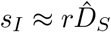 [26, 27].

### Appendix B: QLE Approximation for the Inversion Growth Rate

The general expression in Appendix A depends on the resident disequilibrium 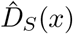, which is generally difficult to obtain for two-locus clines. In the quasi-linkage equilibrium (QLE) regime, recombination is strong enough that disequilibrium is small and can be approximated from single-locus cline shapes. This is the biologically relevant regime for inversion evolution, because if recombination were already weak without an inversion, suppressing recombination would provide little additional benefit.

We assume recombination is fast relative to selection and to spatial turnover on the cline scale (*r* ≫ *s*_0_, *r* ≫ *V*_*m*_*/ω*^2^). For a single locus with allele frequency *p*(*x*), the equilibrium cline satisfies

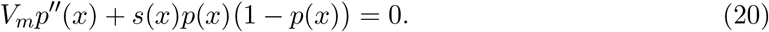

For two loci in QLE, the resident disequilibrium is approximated by

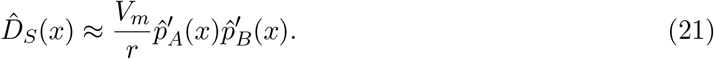

This is the standard statement that linkage disequilibrium is generated by opposing cline slopes and broken down by recombination. Using 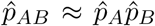 and symmetric loci 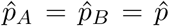, substitution into eq. (17) gives an intermediate expression that still depends on both first and second derivatives of the cline. Taking the limit of strong recombination, *r* → ∞, yields the simpler form

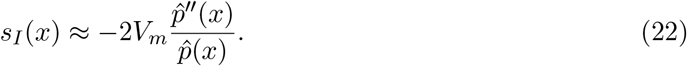

The inversion growth rate is therefore controlled by cline curvature in the high-recombination limit. Rescaling this expression by a characteristic cline width *ω* is carried out in Appendix C.

### Appendix C: Scaling of the Inversion Growth Rate

Appendix B gives the QLE growth rate in physical coordinates,

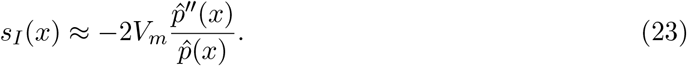

Let *ω* be a characteristic width of the cline and set *χ* = *x/ω*, with 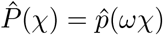. Differentiating with respect to *x* gives

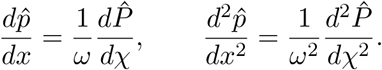

Substituting into the QLE expression yields

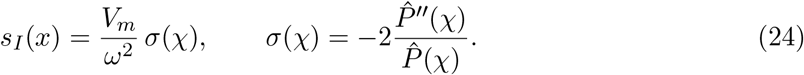

The magnitude of *s*_*I*_ is therefore set by *V*_*m*_*/ω*^2^, while *σ*(*χ*) is a dimensionless spatial profile. When *ω* is chosen so that the scaled cline satisfies eq. (7), 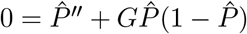 (Appendix D), one has

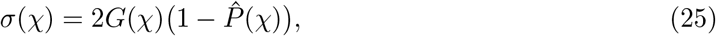

which is eq. (9) of the main text. For each gradient class, *σ* is then independent of *V*_*m*_ and *s*_0_.

### Appendix D: Derivation of Cline-Width Scalings

The main text rescales space by a characteristic width *ω* so that cline shape becomes universal for each environmental gradient. This appendix derives that width and explains how the local inversion growth rate depends on *V*_*m*_, *s*_0_, *L*, and *θ*.

We write selection as *s*(*x*) = *s*_0_*g*(*x*), where *g*(*x*) has transition length *L* and steepness parameter *θ*. Inside the transition region, *g*(*x*) = sgn(*x*)(|*x*|*/L*)^*θ*^, and outside this region *g*(*x*) = sgn(*x*). Substituting the rescaled coordinate *χ* = *x/ω* into the single-locus equation yields

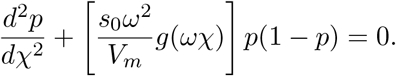

For the equation to become parameter-free for a given gradient class, the prefactor in front of *g*(*ωχ*) must reduce to order one. In a step environment with *θ* = 0, this requires

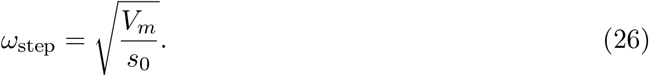

This is the classical cline width [41]. In a linear gradient with *θ* = 1, the selection term contains an explicit factor *χ*, and matching coefficients gives

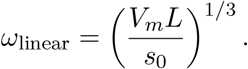

This linear-gradient scaling was first obtained by May et al. [42]. For the general power-law family,

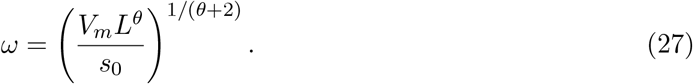

These scalings assume that the environmental transition is wide enough that *L > ω*; if *L* is smaller than the cline width, the system behaves effectively like a step.

Once *ω* is known, the characteristic magnitude of the inversion growth rate is *s*_*I*_ ~ *V*_*m*_*/ω*^2^ (Appendix C). Substituting the general width scaling shows that

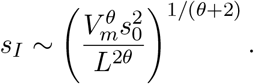

This makes the parameter dependence explicit. For instance, in a step environment with *θ* = 0, *s*_*I*_ can reach *s*_0_ near the center of the cline. For *θ >* 0 ecological tension is relaxed such that the selection differential is spread over space. As a consequence the peak of *s*_*I*_ is lower but the region of positive growth is broader (Fig. 2).

The same expression also implies how sensitivities change with the environmental gradient. For a step environment, increasing *V*_*m*_ does not affect the magnitude of *s*_*I*_. For larger *θ, s*_*I*_ becomes more sensitive to dispersal and less sensitive to selection. For example, doubling *V*_*m*_ at fixed (*s*_0_, *L, θ*) leaves the scaling of *s*_*I*_ unchanged at *θ* = 0, but increases it by a factor 2^1*/*2^ at *θ* = 2.

### Appendix E: Rescaled Establishment Equation

The local growth rate *s*_*I*_ tells us whether a rare inversion grows at a given position, but establishment requires that a single founding lineage survive drift and dispersal long enough to become secure. This appendix derives the establishment probability and explains why it is much lower than the panmictic benchmark and why its maximum is displaced from the cline center.

Consider a rare inversion introduced at position *x* with local growth rate *s*_*I*_(*x*) and dispersal variance *V*_*m*_. During establishment, lineage number is small and genetic drift dominates, so the process can be approximated as a branching process with spatial diffusion. In continuous space this leads to

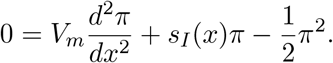

The first term describes dispersal of the founding lineage, the second term describes local growth at rate *s*_*I*_, and the nonlinear term describes the reduction in establishment probability as the lineage becomes common locally.

Because *s*_*I*_(*x*) scales as *V*_*m*_*/ω*^2^ (Appendix C) and *π*(*x*) inherits the same scale, while space scales with *ω*, it is useful to write

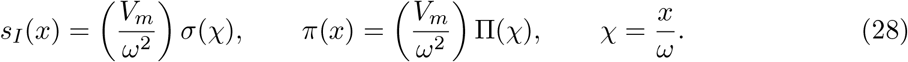

Substituting these definitions and dividing by the common prefactor yields the dimensionless equation

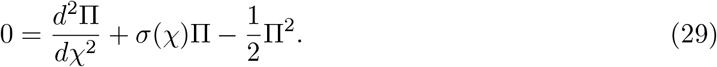

For a fixed environmental gradient, the shape of Π(*χ*) is therefore universal; only its magnitude is set by *V*_*m*_*/ω*^2^.

To interpret the magnitude of Π, it is helpful to compare with a well-mixed population. For a beneficial mutation with constant selection coefficient *s*, classical branching-process theory gives

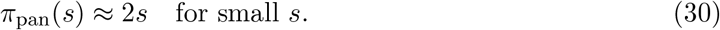

In a step-like cline, the dimensionless growth profile can reach *σ*_max_ ≈ 1, which might suggest naively that Π_max_ should be of order two. Numerical solution of eq. (29) instead gives Π_max_ ≈ 0.1, corresponding to *π*_max_ ~ *s*_0_*/*10 after rescaling. The reduction is not a numerical artifact. It follows because a founding lineage in space does not experience a constant selective environment.

### Appendix F: Total Establishment Rate Scaling

Local establishment probability answers where a single inversion is likely to fix. Population-level questions require the total rate at which successful inversions enter the species. This appendix derives that rate and explains why its dependence on parameters differs from the dependence of the local peak of *s*_*I*_.

The total successful inversion input is

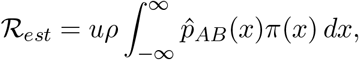

where *u* is the mutation rate and *ρ* is population density. The integrand combines two spatial effects: where adapted haplotypes are common enough to generate inversion mutations, and where those mutations have high establishment probability. Under QLE, 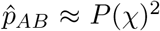, and changing variables to *χ* = *x/ω* gives

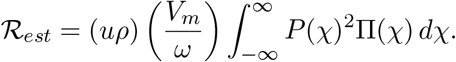

The integral over *χ* depends only on environmental gradient. All dimensional dependence is carried by the prefactor *V*_*m*_*/ω*. For a step environment, ℛ_*est*_ scales as 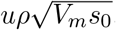, so dispersal and selection contribute with equal strength. For a linear gradient, the scaling becomes 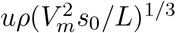. More generally,

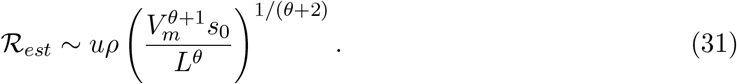

As *θ* increases, ℛ_*est*_ becomes more sensitive to dispersal and less sensitive to peak selection. In practical terms, flatter environmental transitions (*θ >>* 0) can lower the peak of *s*_*I*_ while still increasing the total number of successful inversions, because establishment becomes possible over a wider geographic interval.

### Appendix G: Multi-Locus Generalization under QLE

The main text focuses on two loci, but many inversions capture multiple locally adapted sites. Under QLE, the same logic extends with only modest additional complexity.

Consider an inversion capturing haplotype *H* = *A*_1_*A*_2_ · · · *A*_*n*_. The general two-locus expression still applies, with a total recombination load term and a curvature term for the captured haplotype frequency. Starting from eq. (17), the growth rate is

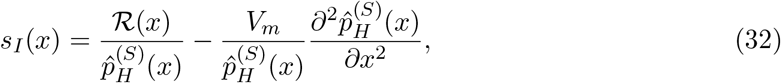

where ℛ(*x*) is the total recombination load breaking down haplotype *H*. Under QLE, we can write the haplotype frequency as a product of single-locus clines plus small LD corrections:

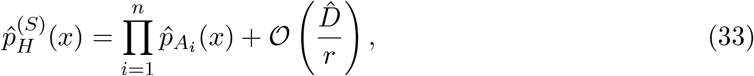

where 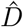 represents pairwise linkage disequilibria, which are O(1*/r*) in the QLE regime. Differentiating the product term gives

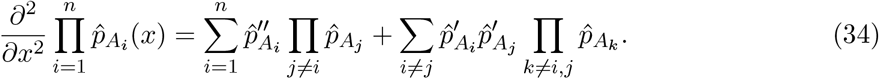

Dividing by 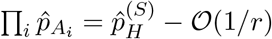 gives

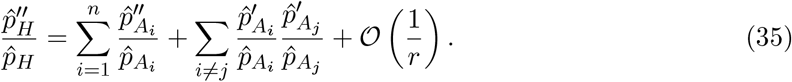

The total recombination load can be approximated in terms of pairwise LDs:

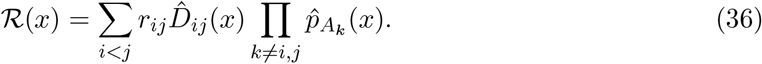

Dividing by 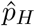 gives

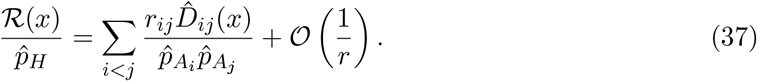

Under QLE, the LD is given by

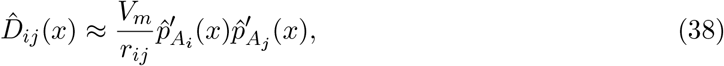

so that

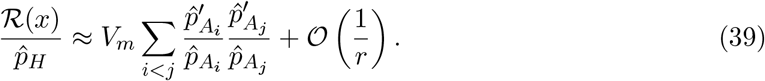

Substituting eq. (35) and eq. (39) into the expression for *s*_*I*_(*x*), we see that the cross-derivative terms cancel exactly:

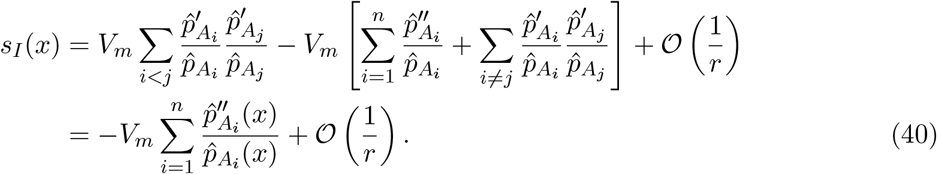

Using the single-locus equilibrium condition,

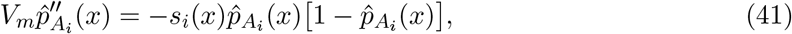

we finally obtain

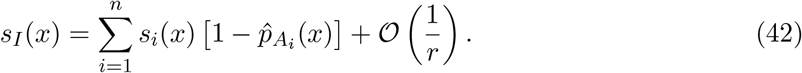

If *s*_*i*_(*x*) = *s*(*x*) and 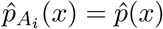 for all *i*, this reduces to

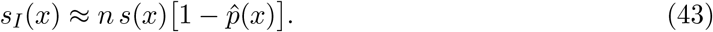

